# A total synthetic approach to CRISPR/Cas9 genome editing and homology directed repair

**DOI:** 10.1101/359984

**Authors:** Sara E. DiNapoli, Raul Martinez-McFaline, Caitlin K. Gribbin, Paul Wrighton, Courtney A. Balgobin, Isabel Nelson, Abigail Leonard, Carolyn R. Maskin, Arkadi Shwartz, Eleanor D. Quenzer, Darya Mailhiot, Clara Kao, Sean C. McConnell, Jill L.O. de Jong, Wolfram Goessling, Yariv Houvras

## Abstract

CRISPR/Cas9 has become a powerful tool for genome editing in zebrafish that permits the rapid generation of loss of function mutations and the knock-in of specific alleles using DNA templates and homology directed repair (HDR). We compared synthetic, chemically modified sgRNAs to in vitro transcribed sgRNAs and demonstrate the increased activity of synthetic sgRNAs in combination with recombinant Cas9 protein. We developed an in vivo genetic assay to measure HDR efficiency and we utilized this assay to optimize the design of synthetic DNA templates to promote HDR. Utilizing these principles, we successfully performed knock-in of fluorophores at multiple genomic loci and demonstrate transmission through the germline at high efficiency. We demonstrate that synthetic HDR templates can be used to knock-in bacterial nitroreductase (*ntr*) to facilitate lineage ablation of specific cell types. Collectively, our data demonstrate the utility of combining synthetic sgRNAs and dsDNA templates to perform homology directed repair and genome editing in vivo.

## INTRODUCTION

CRISPR/Cas9 has been used for a wide range of experimental applications, and zebrafish has been a key model organism to test and validate strategies for genome editing {Jao, 2013 #1}{Shah, 2015 #3}. Repair of CRISPR-generated double-stranded breaks (DSBs) by non-homologous end joining (NHEJ) leads to insertions and deletions (indels) which may result in loss of function of the targeted gene product. By supplying an exogenous DNA template, DSBs can be repaired through homology directed repair (HDR), allowing for precision genome editing, including base pair changes and insertion of protein tags. Prior studies reporting genome editing in zebrafish have used single-stranded donor oligonucleotides (ssODN) to knock-in short DNA sequences {Hruscha, 2013 #34}{Burg, 2018 #35} {Irion, 2014 #7} {Hwang, 2013 #42} or plasmid-based donor vectors to knock-in fluorophores {Auer, 2014 #6}{Hisano, 2015 #36} {Hoshijima, 2016 #9} {Ota, 2016 #38} {Kimura, 2014 #11}. The efficiency of transmitting fluorophore knock-in through the germline has varied widely. We reasoned that the use of synthetic reagents could permit a comparison of approaches and rational optimization.

Reports have described an increased efficiency of CRISPR/Cas9 targeting in human cells using chemically modified synthetic sgRNAs {Hendel, 2015 #12}{Rahdar, 2015 #13}. We evaluated synthetic sgRNAs and found that they outperform conventional in vitro transcribed (IVT) sgRNAs. We used these sgRNAs in combination with synthetic DNA templates and we developed an assay for HDR in zebrafish utilizing an *mitfa* mutant, b692. This assay allowed us to quantitatively compare multiple templates for HDR and correlate phenotypic and molecular efficiency. We performed precise genomic editing and generated in-frame gene fusions with fluorophores using synthetic reagents, including linear dsDNA templates. Knock-in alleles were transmitted through the germline at efficiencies of 14–25%. Finally, we used HDR to target bacterial nitroreductase (*ntr)* to the liver-specific gene *fabp10a* to perform lineage ablation of hepatocytes. These results demonstrate that the combination of synthetic sgRNAs and dsDNA templates result in efficient genome editing in zebrafish.

## MATERIALS AND METHODS

### sgRNA and HDR template sequence selection

Gene-specific sgRNAs sequences were selected using a combination of prediction tools including sgRNA Scorer 1.0 and 2.0{Chari, 2015 #14}{Chari, 2017 #15}, GuideScan{Perez, 2017 #16}, or CRISPRz{Varshney, 2016 #17}. We selected sgRNA sequences with zero predicted off targets with one base pair mismatch. Design principles for HDR templates included a mutated sgRNA recognition sequence, incorporation of barcoded nucleotides, homology arms, and incorporation of heterologous DNA sequence encoding EGFP or mScarlet. HDR templates were ordered as gBlock Gene Fragments (Integrated DNA Technologies [IDT]). For single stranded DNA templates Ultramer DNA Oligonucleotides (IDT) were used. gBlocks were resuspended in nuclease-free water to a concentration of 50 ηg/μL and stored at −20°C.

### Preparation of sgRNAs

Synthesis of IVT sgRNAs was performed from dsDNA templates (gBlock, IDT) using the SureGuide gRNA Synthesis Kit (Agilent, 5190-7719). Templates contained a T7 promoter and GG dinucleotide to promote transcription (TAATACGACTCACTATAGG), a 20bp target site without PAM site, and a composite crRNA/tracrRNA single guide RNA sequence (sgRNA) (GTTTTAGAGCTAGAAATAGCAAGTTAAAATAAGGCTAGTCCGTTATCAACTTGAAAAAGTGGCACCGAGTCGGTGCTTTT). 50ηg of dsDNA template (10ηg/μL) was in vitro transcribed in a 25μL reaction and incubated at 37°C for 3–4 hours. Following DNAse digestion sgRNAs were column-purified and eluted in H_2_O per the manufacturer’s (Agilent) instructions. Concentration was determined using a Nanodrop spectrophotometer and 200ηg of the sgRNA was visualized on an agarose gel for quality control.

Gene-specific crRNAs were synthesized by IDT as Alt-R^®^ CRISPR-Cas9 crRNAs. A bipartite synthetic sgRNA was heteroduplexed using gene-specific crRNAs and a tracrRNA according to manufacturer recommendations. For simplicity the bipartite synthetic sgRNA is referred to as a synthetic sgRNA throughout the manuscript.

### Microinjection

For global editing with a single sgRNA, 250ρg sgRNA and 500ρg recombinant Cas9 protein (rCas9, PNA Bio CP01) were injected into the yolk of one-cell stage embryos. Cas9 nickase (D10A Cas9 nickase protein with NLS, PNABio CN01) was purchased from PNA Bio. To generate deletions, 250ρg sgRNA 1, 250ρg sgRNA 2, and 500ρg rCas9 were injected into the yolk of one-cell stage embryos. For HDR injections, 250ρg sgRNA, 500ρg rCas9, and 37.5ρg HDR template were injected into one-cell stage embryos. Throughout the manuscript rCas9 refers to recombinant Cas9 protein.

### Assessment of gene editing efficiency

Genomic DNA was isolated from individual embryos (24–48 hours post fertilization [hpf]) using DirectPCR Lysis Reagent (Viagen, 102-T) supplemented with Proteinase K at 20μg/mL (Qiagen, 158920). Samples were incubated at 55°C for 60 min followed by 85°C for 45 min. Genomic DNA was PCR amplified using specified primers (Supplementary Table). 8 μL of PCR product was mixed with 1.6μL NEB Buffer 2 and 6.4μL of water and was incubated at 95°C for 5 minutes followed by cooling from 95°C– 85°C at −2°C/second and 85–25°C at −0.1°C/second. Following hybridization, the DNA was subject to T7 endonuclease (T7EI) digestion with 2U of T7EI endonuclease (New England BioLabs [NEB], M0302) and 1 hour incubation at 37°C. The reaction product was visualized on a 2% agarose gel.

For clonal analysis, the PCR product was cloned into pCRII-TOPO (Invitrogen, K460001) and Sanger sequenced (Genewiz). Sequences were compared to reference genome and non-targeting controls to identify indels using CrispRVariants{Lindsay, 2016 #18} and MacVector (Version 15.5).

For CRISPR-STAT analysis, sperm samples from potential founders were collected with 10μL capillary tubes as previously described, and then followed the HOTShot method with 25μL alkaline lysis buffer to prepare DNA templates{Draper, 2009 #19}{Meeker, 2007 #20}. Tail clips were prepared similarly. The CRISPR-STAT protocol was used to perform fluorescence-based genotyping{Carrington, 2015 #21}. Primers sequences used for the CRISPR-STAT assay were *slc6a15* FOR, tgtaaaacgacggccagtGGCCACGACCTACTACTGGTAT, which includes (lowercase) M13 tag for binding to a fluorescent-labeled M13 primer; and *slc6a15* REV, gtgtcttTATAGATGCTGCGTCACGTTTC, which include the (lowercase) ‘PIG tail’ tag that helps ensure uniform product size, via Taq adenylation{Holleley, 2009 #22}. Relative amounts of each uniquely sized PCR product (>100 bp and >100 peak height) were calculated with area under the curve measurements using Applied Biosystems Peak Scanner software.

For assessment of deletions, genomic DNA was PCR amplified using primers that flanked the deletion sgRNAs. PCR products were purified (Qiagen, 28104), Sanger sequenced, and compared to the reference genome.

### Analysis of HDR efficiency by next-generation sequencing

For analysis of HDR efficiency by next-generation DNA sequencing, 50ηg of PCR amplicon was converted into blunt ends using T4 DNA polymerase (New England Biolabs, NEB) and *E. coli* DNA polymerase I Klenow fragment (NEB). Libraries were prepared using the Kapa LTP Library Preparation Kit (Roche). Multiple indexing adapters were ligated to the ends of the DNA fragments. Ligation reaction products were purified by AMPure XP beads (NEB) to remove unligated adapters and quantified using Qubit (Thermo Fisher Scientific) and Bioanalyzer DNA chip (Agilent). Indexed sample libraries were normalized, pooled, and sequenced using the Illumina HiSeq4000 sequencer at 2×50 cycles. Reads were aligned to the zebrafish genome (GRCz10) using the Star aligner{Dobin, 2013 #23} and visualized using Integrative Genome Browser (IGV) {Thorvaldsdottir, 2013 #24}{Robinson, 2011 #25}.

### Imaging

Micrographs of whole embryos and larval animals were taken with a Zeiss Discovery V8 stereomicroscope (Zeiss) equipped with epifluorescence and appropriate filters. Live imaging of fluorescent larvae was acquired with a Zeiss LSM800 laser scanning confocal microscope (Zeiss). For melanocyte counting, 48hpf embryos were visualized under the stereoscope and binned into 4 categories (WT, >101, 51–100, or 0–50 melanocytes). Individuals scoring embryos were blinded to the experimental conditions.

### Hepatocyte ablation

HDR injection mixes were made as described above. Approximately 2 ηL of mix was injected into the cell of one-cell stage wild-type (Tu) embryos. At 4–6 days post fertilization (dpf), embryos were screened by confocal microscopy for mosaic mScarlet expression localized to the liver, and positive larvae were raised to adulthood. F_0_ adults were outcrossed with wild-type (TL) fish, and the F_1_ embryos were screened by confocal microscopy for mScarlet positive livers. Positive larvae were divided and treated with either 0.2% dimethylsufoxide (DMSO; Sigma Aldrich) or 10 µM metronidazole (Mtz; Sigma Aldrich 443-48-1) in egg water. After 24 hours, the larvae were washed, anesthetized with 0.16 mg/mL tricaine-S (MS 222; Western Chemical, Inc.), and immobilized in 0.8% low melt point agarose (Invitrogen) on glass-bottom dishes (MatTek). Imaging was performed using a Nikon Ti2 inverted microscope equipped with a Yokogawa CSU-W1 spinning disk confocal unit and a Zyla 4.2 PLUS sCMOS camera (Andor), using CFI Plan Apochromat Lambda 10x NA 0.45 and 20x NA 0.8 objectives. Larvae were recovered from the agarose, allowed to recover in the absence of drug for 48 hours, and imaged again.

### Zebrafish husbandry

Zebrafish strains were maintained according to established guidelines{Westerfield, 2007 #33}. Wild-type animals were from the AB background except when indicated. Experiments were performed in accordance with the recommendations in the Guide for the Care and Use of Laboratory Animals of the National Institutes of Health. All of the animals were handled according to approved institutional animal care and use committee (IACUC) protocols of the respective institutions.

## RESULTS

### Synthetic sgRNAs lead to highly efficient induction of indels

In order to directly compare editing efficiencies generated by synthetic sgRNA versus IVT sgRNA), we targeted tyrosinase (*tyr*), a gene required for melanin synthesis. Wild-type zebrafish embryos were microinjected at the one-cell stage with recombinant Cas9 protein (rCas9) complexed with either synthetic sgRNA (IDT, Alt-R^®^) or with laboratory-synthesized IVT sgRNA. Embryos were scored for melanocyte number at 48 hours post fertilization (hpf) and divided into phenotypic categories representing the degree of gene editing (Figure 1A). We find that the synthetic sgRNA leads to a significantly increased fraction of embryos with fewer melanocytes than the IVT sgRNA (Figure 1B). Higher doses of synthetic sgRNA produce an increase in the fraction of phenotypically edited embryos. Only synthetic sgRNAs were capable of producing large numbers of embryos with high phenotype. In order to characterize the molecular alterations of synthetic sgRNA injected embryos, we performed a clonal analysis. We find that 10/10 clones from a synthetic sgRNA injected high-phenotype embryo contain indels, as compared with 7/10 clones from an IVT sgRNA injected high-phenotype embryo; we also observe significant differences in the indel spectrum (Figure 1C). We found comparable differences between synthetic and IVT sgRNAs at two additional loci, *slc24a5* and *slc45a2* (Supplementary Figure 1). Injection of synthetic sgRNAs leads to highly penetrant phenotypes in F_0_ larvae that develop into adults that phenocopy established germline mutants{Sheets, 2007 #26} (Supplementary Figure 2). We find that pairs of synthetic sgRNAs are able to induce deletions spanning up to 124kB in genomic DNA and that the efficiency of generating deletions varies with size (Supplementary Figure 3). Taken together, these data indicate that combined use of rCas9 protein with synthetic sgRNAs leads to penetrant phenotypes and a broad spectrum of gene-specific edits.

**Figure 1.**
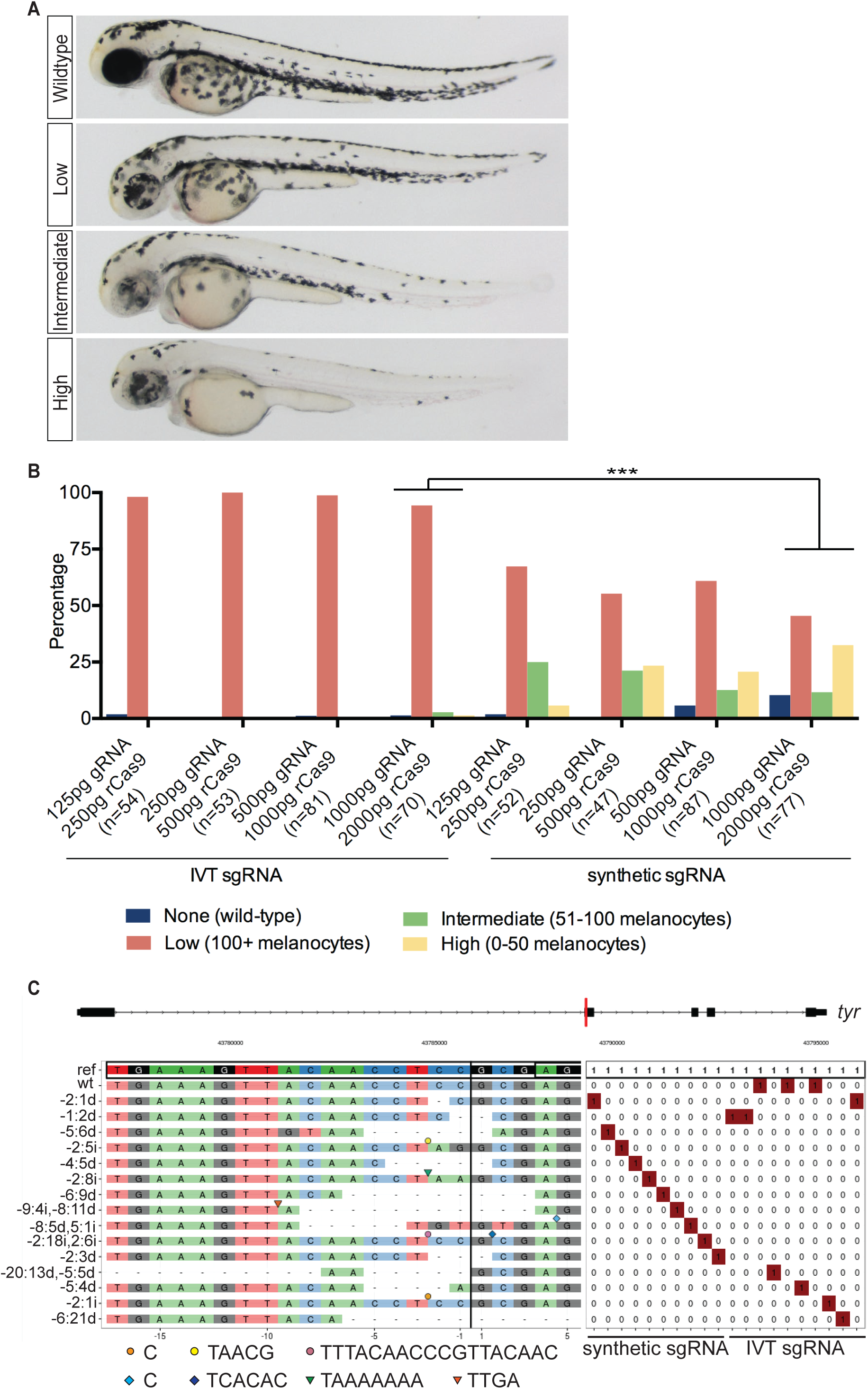
Synthetic sgRNAs outperform laboratory synthesized *in vitro* transcribed sgRNAs. (**A**) Dose-response comparison of editing in embryos injected with rCas9 and either an *in vitro* synthesized or synthetic sgRNA targeting *tyr*. Embryos were scored for melanocyte dropout at 48hpf and binned into 4 categories. The percentage of embryos in each condition is plotted. At the 1000pg sgRNA/2000pg rCas9 dose, there is a statistically significant increase in the editing observed with the synthetic sgRNA (p=<0.001, Chi-square). (**B**) Light microscopic images of representative embryos in each category. (**C)** CrisprVariants {Lindsay, 2016 #18} plot of indels in clones isolated from embryos injected with either synthetic or *in vitro* transcribed sgRNAs. 10/10 clones isolated from individual embryos injected with synthetic from sgRNAs contained indels and 7/10 clones isolated from embryos injected with IVT sgRNAs contained indels.

To assess the extent of germline transmission of edited alleles, we applied CRISPR-STAT fluorescence-based genotyping to F_0_ founders and progeny{Carrington, 2015 #21}. Sperm samples were collected from F_0_ founders injected with rCas9 and synthetic sgRNA targeting *slc6a15* at the one-cell stage. Results reveal mutant alleles containing indels as the major peaks, with additional minor peaks corresponding to wild-type and mutant alleles, indicating a high efficiency of CRISPR-induced mutagenesis in the germline (Supplementary Figure 4). We identified five distinct alleles that were transmitted to F_1_ progeny (Supplementary Figure 4B-F). The frequencies of observed CRISPR-induced indels in F_1_ progeny were similar to those found in the sperm of the F_0_ founder (Supplementary Figure 4G). These findings are consistent with high levels of somatic mosaicism and confirm that F_0_ sperm samples accurately predict the specific mutations transmitted to F_1_ progeny. These data demonstrate that CRISPR-induced indels created with synthetic sgRNAs are found in the germline of founders and are efficiently transmitted to F_1_ progeny.

### A genetic assay for homology-directed repair

To develop a quantitative assay for optimizing HDR in zebrafish, we sought to target a locus that could provide phenotypic and molecular data on allele conversion. We initially examined the widely used *mitfa*(w2) mutant, which lacks melanocytes{Lister, 1999 #27}. Unexpectedly, injection of rCas9 complexed with a synthetic sgRNA targeting the mutation site in w2 resulted in restoration of pigmented melanocytes (Supplementary Figure 5). We presume that indels caused by targeting the *mitfa*(w2) mutation site (Q113Stop) lead to in-frame deletions and restore protein function since this region of the protein is relatively unstructured. We next examined the *mitfa*(b692) mutant, which lacks melanocytes and harbors a mutation leading to a Ile215Ser substitution{Lister, 2001 #28}. We find that a synthetic sgRNA targeting the *mitfa*(b692) mutation site does not restore pigmented melanocytes (Supplementary Figure 5). Therefore, we used *mitfa*(b692) to develop a quantitative phenotypic and molecular assay for optimizing HDR.

Using the b692-HDR assay we examined a range of synthetic templates to optimize HDR efficiency. *mitfa*(b692) embryos were microinjected with rCas9, synthetic sgRNA, and synthetic DNA templates designed to restore the wild-type *mitfa* sequence and gene function. Microinjected embryos were scored at 48hpf for phenotypic evidence of melanocytes, indicative of allele conversion and mitfa function (Figure 2A). Melanocytes were not identified in uninjected embryos or in embryos injected with non-targeting sgRNA. HDR templates were designed to feature multiple barcoded nucleotides and mutated sgRNA recognition sequence, enabling unambiguous detection by sequencing and prevention of re-cleavage by rCas9. We found that a 951bp, double-stranded linear DNA template with an asymmetric sgRNA site located at the 3’ end leads to a reproducible 9% rate of phenotypic rescue in *mitfa*(b692) embryos (Figure 2B). Melanocyte-positive embryos were subject to allele-specific PCR and Sanger sequencing to confirm precise HDR (Figure 2C-D).

**Figure 2.**
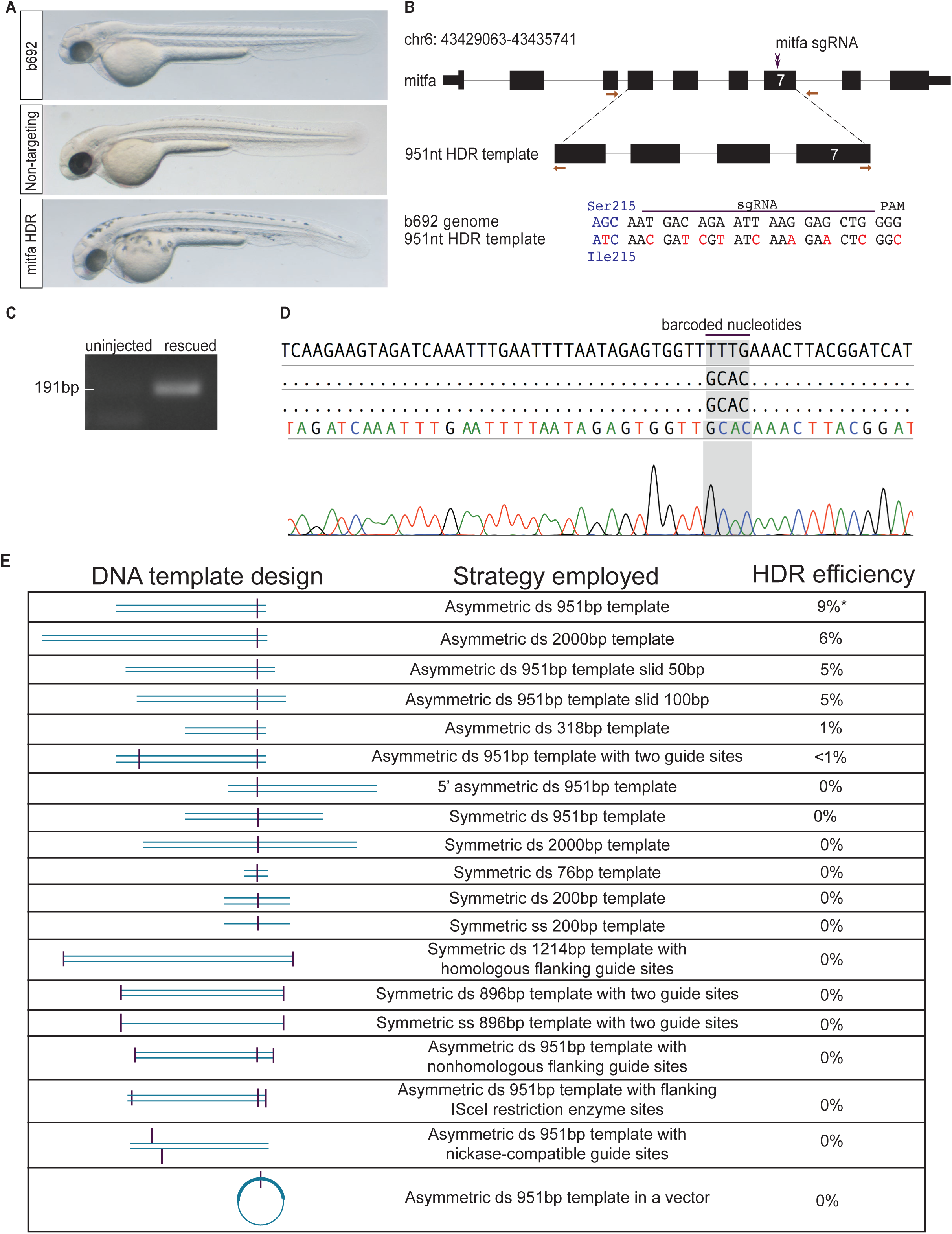
A genetic assay for optimizing homology-directed repair using b692 mutant zebrafish. (**A**) *mitfa*(b692) embryos uninjected (top), injected with rCas9, 951bp DNA template, and non-targeting sgRNA (middle) or *mitfa* sgRNA (bottom). All embryos shown at 48hpf. (**B**) Schematic depicting *mitfa* gene structure and location of the mutation in exon 7 in b692 mutants. The sgRNA location is indicated. Primers used for allele-specific PCR are displayed as orange arrows. A linear 951bp dsDNA template encodes a wild-type Ile codon at position 215 and additional nucleotide changes to prevent re-cleavage at the sgRNA recognition site (red). (**C**) Allele-specific PCR was performed on rescued embryos after microinjection with 951bp template and compared with uninjected embryos. (**D**) Sanger sequencing results of allele-specific PCR products from a phenotypically rescued embryo. The sequence read spans the template-genome junction and includes template specific barcoded nucleotides. (**E**) Chart depicting attributes of DNA template (left), a description of the template (middle) and observed rate of rescue in the b692-HDR assay (right).

We examined DNA templates that differ in length, single vs. double stranded DNA, linear vs. circular templates, and the symmetry of the sgRNA position within the template (Figure 2E). We find that both longer (2kB) and shorter (318bp, 76bp) DNA templates led to a decrease in HDR efficiency. Asymmetric positioning of sgRNA sites led to higher efficiency than symmetric positioning. We tested an IVT sgRNA in combination with rCas9 and the optimal 951bp linear dsDNA template and found no appreciable HDR. Our data indicates that HDR efficiency is maximal with dsDNA templates and synthetic sgRNA and that HDR is sensitive to template length and sgRNA site symmetry within the HDR template.

Using the b692 assay we examined multiple reagents and conditions reported to improve HDR efficiency in other model systems (Figure 2E). The addition of a second sgRNA site in the optimal 951bp template reduced HDR efficiency to < 1%. We cloned the 951bp template into a TopoTA plasmid vector and microinjected a circular plasmid into zebrafish. This plasmid-based template also failed to lead to measurable HDR in b692 embryos. In light of prior studies reporting increased HDR efficiency with the use of catalytically-mutated Cas9 (nickase Cas9) {Richardson, 2016 #29}, we altered the 951bp DNA template to allow us to test this approach. Microinjection of paired sgRNAs and recombinant nickase Cas9 failed to lead to measurable HDR in b692 embryos. Based on reports that small molecule inhibitors of poly (ADP-ribose) polymerase (PARP) stimulate HDR efficiency {Anantha, 2017 #32}, we performed microinjection of the 951bp template and a PARP inhibitor, but did not observe any measurable HDR efficiency. Finally, we examined the effect of cleaving the ends of the linear 951bp template with a restriction enzyme (ISce-I), but we did not observe measurable HDR under this condition.

### Melanocyte restoration in b692 assay correlates with molecular efficiency

Using the b692-HDR assay we sought to measure the molecular efficiency of genome editing and correlate DNA template integration with melanocyte number. To this end, we performed single-cell injections in a large cohort of b692 embryos using the optimized reagents described: rCas9, synthetic b692 sgRNA, and 951bp linear dsDNA template. Embryos were scored for the presence of melanocytes at 48hpf and binned by phenotype into four categories: high-, medium-, low-, and no-editing. Uninjected embryos were included as a negative control. Genomic DNA was extracted from pools of five embryos in each category, and exon 7 was amplified by PCR. Amplicons from each group were subjected to next-generation sequencing. We find that that the efficiency of genome editing as determined by next-generation sequencing correlates with phenotype in b692 edited embryos. In high-rescued embryos, we find that 17% of reads (106428 edited/641281 total) harbor an edited codon restoring Ser215 to Ile, consistent with template integration. Analysis of aligned reads at exon 7 reveals the uniform presence of barcoded nucleotides from the HDR template across the sgRNA target site (Figure 3A). Phenotypic rescue in medium and low categories revealed molecular efficiency of approximately 3% (14810/575344 and 17566/622068 reads, respectively). Injected embryos with no phenotypic evidence of HDR had molecular efficiency of <0.3%. Alignment of reads demonstrates a mutually exclusive pattern of either HDR with template integration or indels at the sgRNA site (Figure 3A), consistent with highly efficient dsDNA cleavage. We analyzed the molecular efficiency across all phenotypes and plotted the fraction of reads harboring barcoded nucleotides from the HDR template (Figure 3B-E). These data confirm the sensitivity of the b692-HDR assay to read out the molecular efficiency of HDR in vivo.

**Figure 3.**
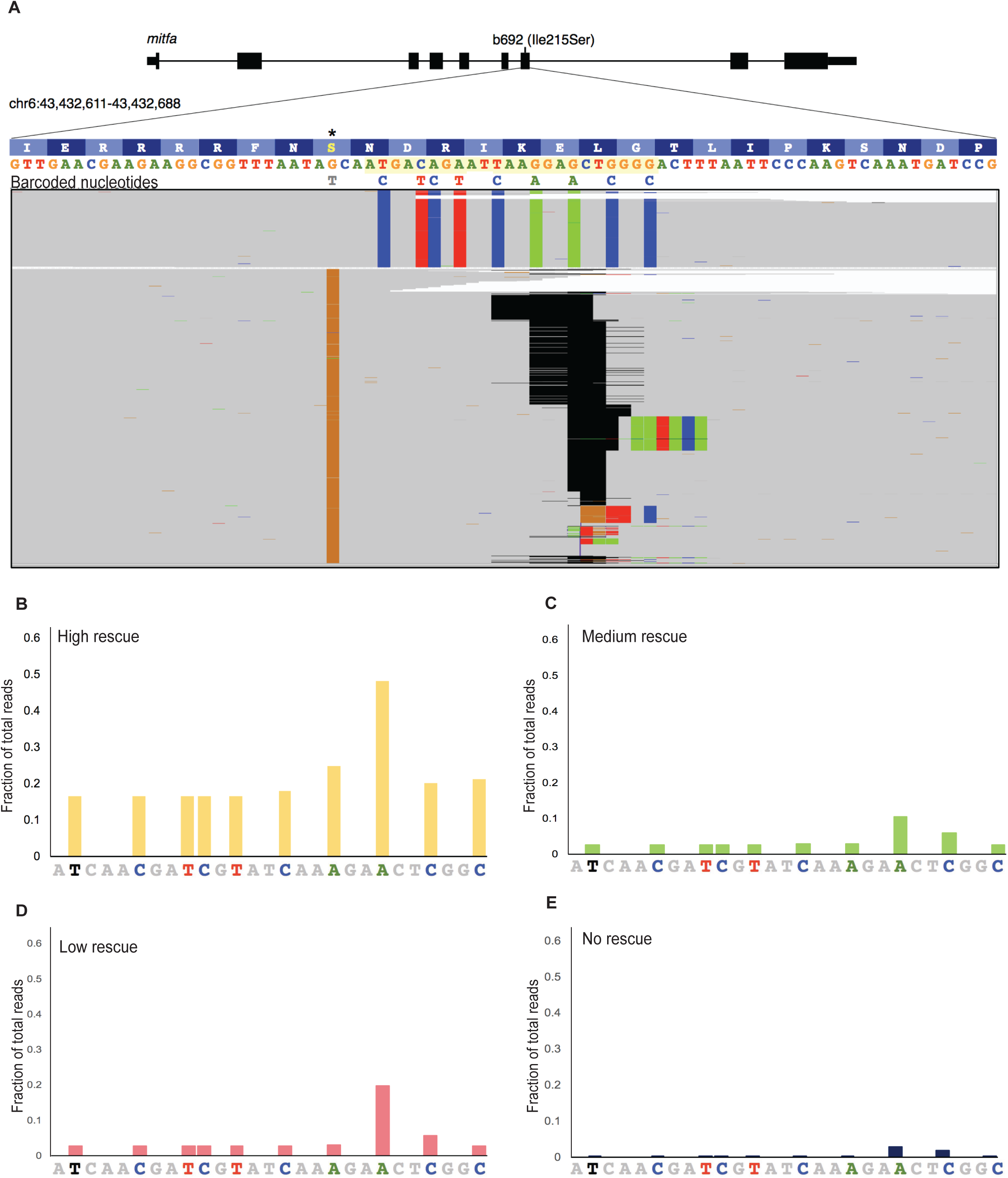
Quantitative assessment of genome editing efficiency by next-generation sequencing. (**A**) Alignment of next generation sequencing reads from b692 genome edited embryos from the high rescue phenotype. Exon 7 of *mitfa* is displayed. The sgRNA sequence is highlighted in yellow. b692 mutants have a T>G mutation leading to an isoleucine to serine substitution at codon 215(*). The 951bp dsDNA template restores the wild-type isoleucine codon (ATC) and encodes nine additional barcoded nucleotides as indicated. Indels in sequencing reads are represented as gapped regions in black. (**B-E**) The fraction of reads at each barcoded nucleotide in the HDR template is plotted for each phenotype category.

### Genome editing using synthetic reagents leads to efficient fluorophore knock-in and germline transmission

To determine the feasibility of using synthetic reagents to knock-in a fluorophore by HDR, we targeted *tyrp1b*, a melanocyte-specific gene. Using albino embryos (pigmentless due to a null mutation in *slc45a2*) we performed microinjection of rCas9 with a synthetic sgRNA and a synthetic linear dsDNA template designed to knock EGFP in frame with the terminal *tyrp1b* exon (Figure 4A). We identified 7/153 (5%) embryos with EGFP-positive melanocytes at 48hpf (Figure 4B). Allele-specific PCR was performed from injected embryos with EGFP-positive melanocytes (Figure 4C). Sanger sequencing revealed the presence of multiple barcoded nucleotides encoded by the HDR template, consistent with template integration (Figures 4D). Confocal imaging revealed the presence of an EGFP-positive melanocyte with characteristic stellate morphology (Figure 4D).

**Figure 4.**
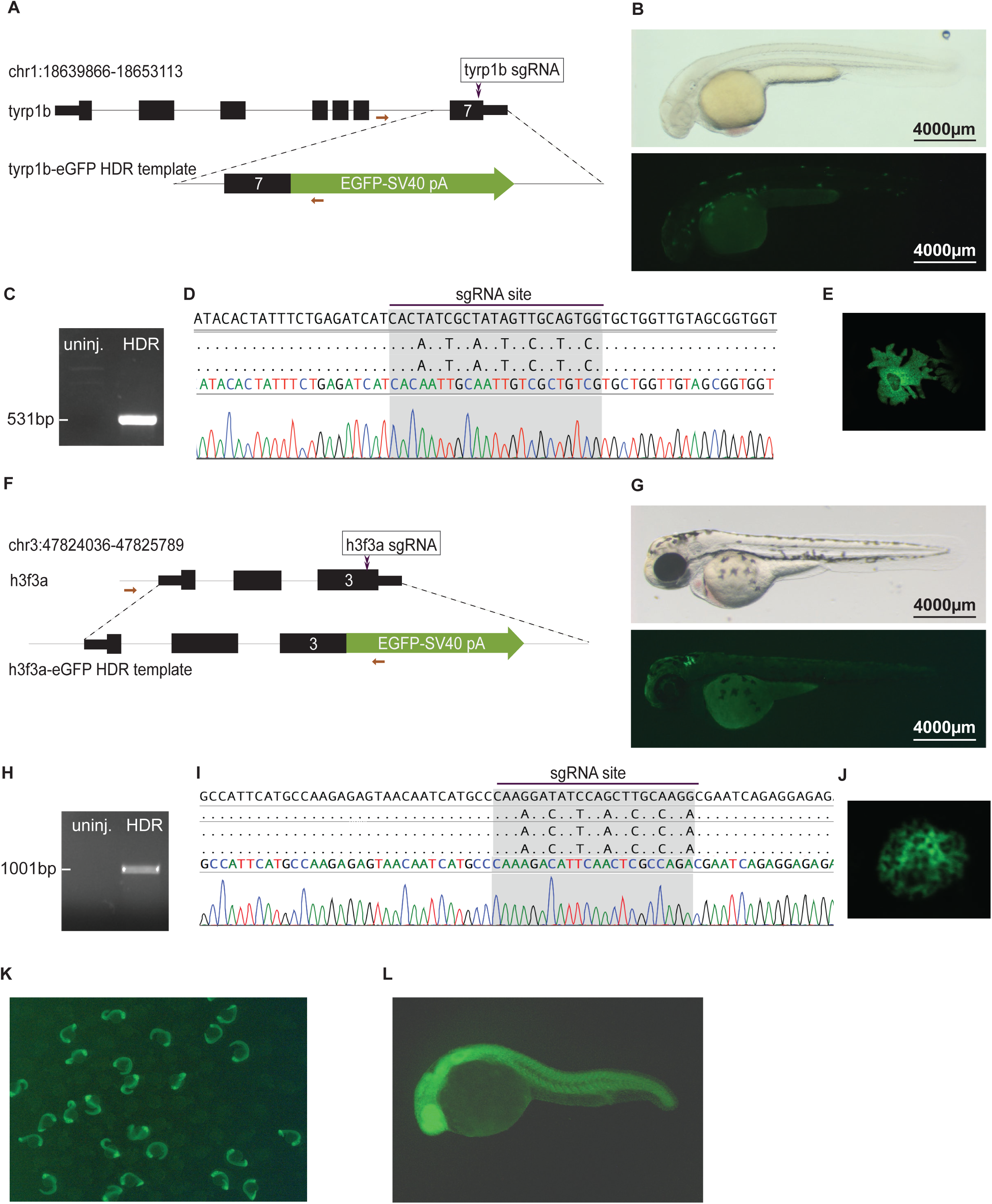
Genome editing leads to precise fluorophore knock-in. (**A**) Schematic depicting *tyrp1b* locus. A linear, dsDNA template designed to lead to an in-frame EGFP knock-in is shown. (**B**) albino(b4) embryo injected with rCas9, *tyrp1b* synthetic sgRNA, and dsDNA template is photographed with light (top panel) or fluorescence (bottom panel) microscopy at 48hpf. (**C**) Allele-specific PCR was performed on uninjected or HDR edited/GFP+ embryos at 48hpf. Only HDR-edited/EGFP-positive embryos had a PCR amplicon. (**D**) Sanger sequencing from allele-specific PCR to detect template integration. The sequence was aligned to reference (top). Barcoded nucleotides at the sgRNA binding site consistent with template integration were identified. A representative chromatogram is shown. (**E**) Confocal imaging reveals the presence of stellate EGFP-positive melanocytes. (**F**) Schematic depicting *h3f3a* locus. A linear, dsDNA template designed to lead to an in-frame EGFP knock-in is shown. (**G**) Wild-type embryo injected with rCas9, *h3f3a* sgRNA, and dsDNA template is photographed with light (top panel) or fluorescence (bottom panel) microscopy (48hpf). (**H**) Allele-specific PCR was performed on uninjected or HDR edited/GFP+ embryos at 48hpf. Only HDR-edited/EGFP-positive embryos had a PCR amplicon. (**I**) Sanger sequencing was performed from allele-specific PCR to detect template integration. The sequence was aligned to reference (top). Barcoded nucleotides at the sgRNA binding site, consistent with template integration were identified. A representative chromatogram is shown. (**J**) Confocal imaging reveals the presence of an EGFP-positive nuclei. (K-L) Immunofluorescent images of F1 progeny showing transmission of EGFP from h3f3a-EGFP founder.

To determine the generalizability of this approach across multiple loci, we targeted *h3f3a*, one of several genes encoding histone H3.3. We designed a linear dsDNA template to knock EGFP in frame with *h3f3a*, creating a C-terminal fusion (Figure 4F). We microinjected rCas9 with a synthetic sgRNA targeting *h3f3a* and a synthetic dsDNA template into wild type embryos. We identified 12/152 (8%) embryos with EGFP-positive cells at 48hpf (Figure 4G). Allele-specific PCR and Sanger sequencing performed on EGFP positive embryos revealed precise template integration (Figure 4H,I). Confocal imaging revealed the presence of cells with a nuclear EGFP-positive signal characteristic of chromatin (Figure 6J). We raised mosaic F_0_ embryos to adulthood and assayed the rate of germline transmission. Of the 14 adult fish that mated, 2 (14%) produced EGFP+ F_1_ embryos (Figure 4K,L). Taken together, these data demonstrate the ability of synthetic sgRNAs and dsDNA templates to knock-in allele-specific fluorophores at several genomic loci and to transmit these edited alleles through the germline.

To expand the applications of genome editing using synthetic DNA templates, we targeted the liver-specific gene *fabp10a* using a linear dsDNA template that knocks in mScarlet and bacterial nitroreductase (*ntr*) to facilitate hepatocyte ablation studies (Figure 5A). The zebrafish liver is capable of regeneration after injury {Cox, 2015 #40}, and expression of *ntr* has been used in combination with metronidazole (Mtz) to cause cellular injury {Curado, 2008 #41}. We injected embryos with rCas9, sgRNA targeting *fabp10a*, and the HDR template and raised mScarlet+ F_0_ embryos to adulthood. Of adult fish that mated, 2/8 (25%) produced mScarlet+ F_1_ embryos. We characterized the recovered *fabp10a* allele in F_1_ animals using allele-specific PCR and confirmed integration of the template in mScarlet+ embryos (Figure 5B). Molecular analysis indicated that the 5’ end of the HDR template resulted in an in-frame fusion between exon 4 and mScarlet. At the 3’ end of the recovered allele, we identified a fragment of exon 4 distal to the sgRNA recognition sequence, likely resulting from ligation to the template. We treated transgenic F_1_ embryos with Mtz to examine whether an fab10a-ntr fusion was functional and could leave to liver-specific cytotoxicity. Exposing fabp10a-mScarlet-NTR larvae to Mtz from 96–120 hpf resulted in a dramatic reduction in the mScarlet fluorescent signal, signifying hepatocyte ablation without obvious damage to other tissues (Figure 5C,D). Remnants of the ablated hepatocytes were observed circulating in the vasculature. After a recovery period of 48 hours in the absence of Mtz, mScarlet signal was observed in the liver of the Mtz-treated larvae, consistent with liver regeneration (Figure 5E,F).

**Figure 5.**
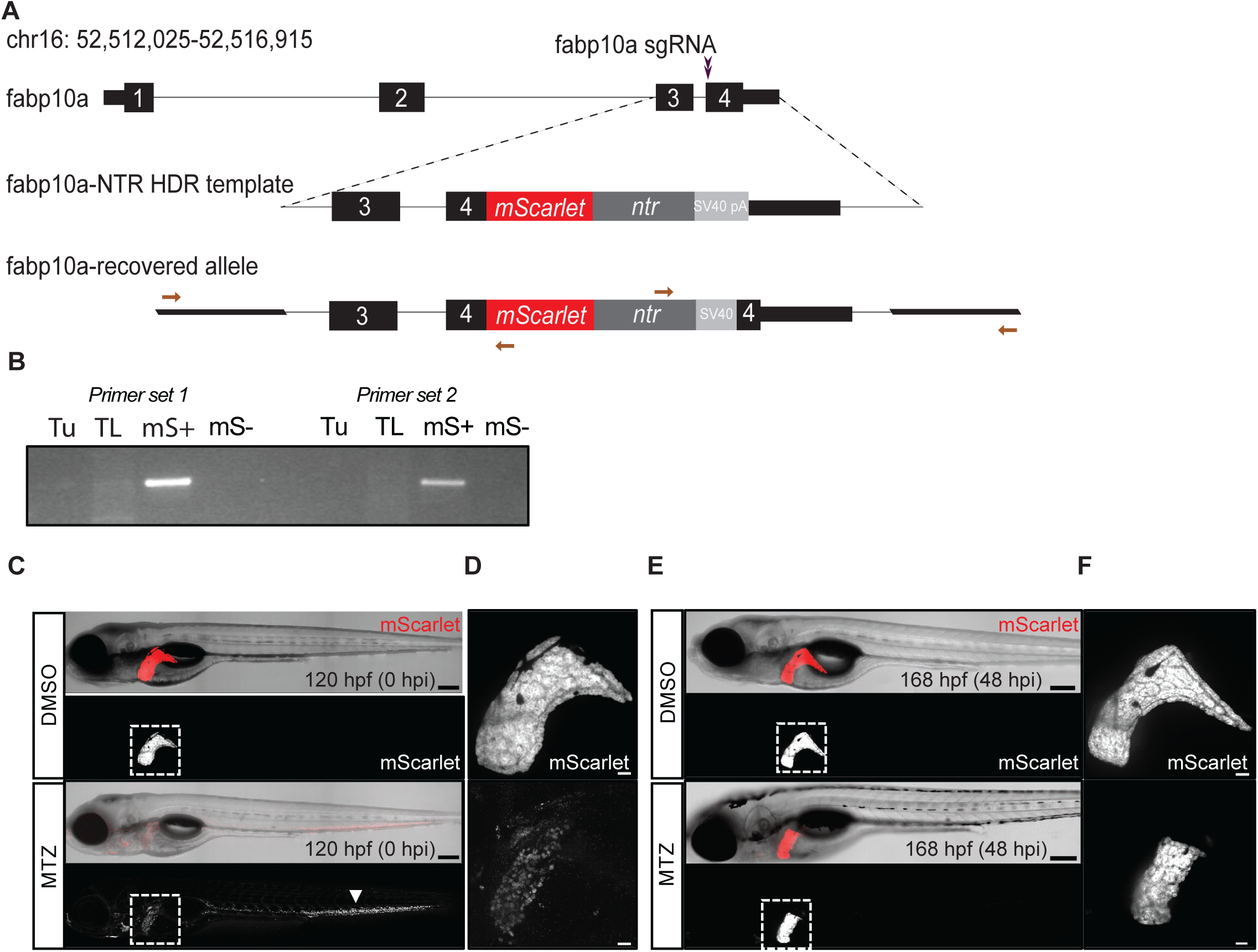
Knock-in of a synthetic dsDNA template encoding a fluorophore and bacterial nitroreductase enables tissue-specific ablation. (**A**) Schematic outlining HDR template design at the *fabp10a* locus. A 2.0kB template was designed to insert mScarlet-NTR as a C-terminal fusion to the *fabp10a* gene, which has hepatocyte-specific expression. The recovered allele from F_1_ progeny confirms integration of mScarlet after the last coding exon of *fabp10a*. **(B)** Allele-specific PCR was performed to confirm integration of the template in mScarlet+ (mS+) embryos. (**C-F**) Mtz exposure induced hepatocyte injury in fabp10a-mScarlet-NTR larvae. **(C)** Confocal imaging revealed loss of fluorescence in the liver (outlined) of Mtz-treated but not DMSO-treated larvae. Remnants of ablated hepatocytes were found distributed throughout the vasculature (arrowhead). **(D)** Confocal imaging of liver in DMSO-and Mtz-treated larvae after injury. (**E**) At 48 hours post injury (hpi), mScarlet signal was observed in the liver (outlined) of the Mtz-treated larvae, demonstrating liver regeneration. **(F)** Confocal imaging of liver in DMSO and Mtz treated larvae at 48 hpi. Scale bars are 200 μm (C,E) and 30 μm (D,F).

## DISCUSSION

Here we describe the development and application of genome editing in zebrafish using synthetic reagents. We demonstrate that the combined use of synthetic sgRNAs with rCas9 protein leads to highly efficient indels. The improved efficiency of commercial sgRNAs may result from chemical modifications that protect sgRNAs from degradation. In order to optimize the efficiency of homology-directed repair we developed a genetic assay using *mitfa*(b692) mutant zebrafish. Our data demonstrate that the b692-HDR assay permits correlation between phenotype and HDR allele conversion (genotype). Using this assay, we systematically tested multiple template designs to determine which parameters are optimal for efficient knock-in by HDR. We find that the optimal template is a linear, dsDNA template with an asymmetric sgRNA site. Using these design principles, we knock-in fluorophores at *tyrp1b, h3f3a* and *fabp10a*; we achieved germline transmission rates of 14–25% at two loci. At *fabp10a* the HDR template included an *ntr* cassette to facilitate lineage ablation studies, and application of metronidazole resulted in hepatocyte ablation in transgenic F_1_ larvae. These germline transmission rates are among the highest reported for knock-in of gene cassettes in zebrafish, and this is the first report of CRISPR-mediated *ntr* knock-in and lineage ablation.

A total synthetic approach has the unique advantage that sgRNA and HDR templates can be designed in silico and commercially manufactured in a short time frame. Design and precision genome editing with a synthetic dsDNA template can be performed within 2–3 weeks. The use of synthetic reagents may reduce experimental variability between labs. The use of sgRNAs produced by commercial solid phase synthesis raises the possibility of incorporating degeneracy in the sgRNA sequence to permit a single sgRNA to target multiple related genes. The synthesis of linear dsDNA templates may allow investigators to rapidly test sequence requirements for HDR. We unexpectedly found significant differences in HDR efficiency between templates with minor design differences. These observations suggest that empiric optimization of donor templates may be required to achieve knock-in at some loci. As new recombinant Cas9 proteins become available, the ability to perform base editing and other enzyme modifications may be possible. Finally, the b692-HDR assay may be used as a screening platform to identify chemical and genetic modifiers of HDR efficiency to make further improvements in efficiency.

The use of CRISPR/Cas9 to generate loss of function mutations in zebrafish and other model organisms makes it a uniquely valuable resource for forward genetics. To our knowledge, this is the first study to develop a genetic assay for HDR in zebrafish and to optimize HDR templates. Our studies offer insight into the relative efficiency of HDR and provide investigators with a workflow for generating knock-in alleles. Our approach to genome editing should allow investigators to pursue a broad range of in vivo applications, including tissue-restricted lineage ablation, fluorophore and epitope knock-in, and generation of conditional alleles. Future improvements in HDR efficiency may occur with new variations in Cas9 protein and molecular and chemical tools to further enhance the efficiency of homology directed repair.

## ACKNOWLEDGMENT

We are grateful to members of the Houvras laboratory for critical discussion and manuscript review. We thank the Weill Cornell Genomics Resources Core Facility for next generation sequencing.

---

**Supplementary Figure 1.**
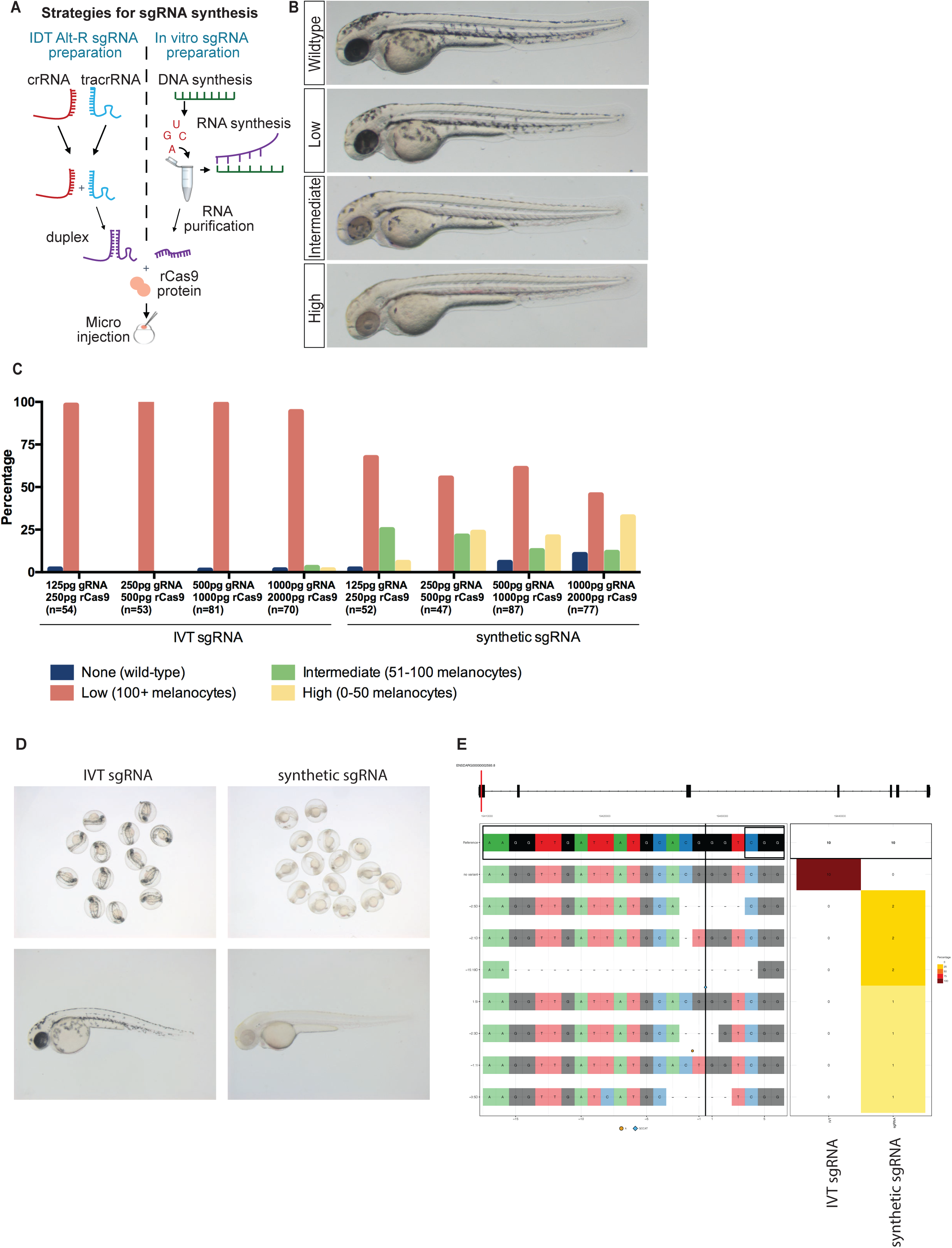
Synthetic sgRNAs outperform *in vitro* synthesized sgRNA in targeting *slc25a2* (golden) and *slc45a2* (albino). (**A**) Comparison of Alt-R^®^ synthetic sgRNA preparation and *in vitro* synthesis of sgRNAs. Left, Alt-R^®^ synthetic sgRNAs are prepared by duplexing a gene specific crRNA with a common tracrRNA. Right, IVT sgRNAs are transcribed from a DNA template using RNA polymerase and purified. Both sgRNAs are used with rCas9 and microinjected into embryos at the one-cell stage. (**B**) Light microscopic images of representative embryos in each phenotypic category. (**C**) Dose-response comparison of editing in embryos injected with rCas9 and either an *in vitro* synthesized or synthetic sgRNA targeting *slc25a2*. Embryos were scored for melanocyte dropout at 48hpf and binned into 4 categories. The percentage of embryos in each condition is plotted. (D) Embryos injected with either IVT or synthetic sgRNA targeting slc45a2 (albino). Representative embryos are shown. IVT sgRNA was ineffective in inducing a mutant phenotype. Synthetic sgRNA induced a mutant phenotype in virtually all injected embryos. (E) A clonal analysis was performed and analyzed using CrisprVariants. IVT injected embryos showed no evidence of indels (0/10 clones) and a synthetic sgRNA induced indels in 10/10 clones.

**Supplementary Figure 2.**
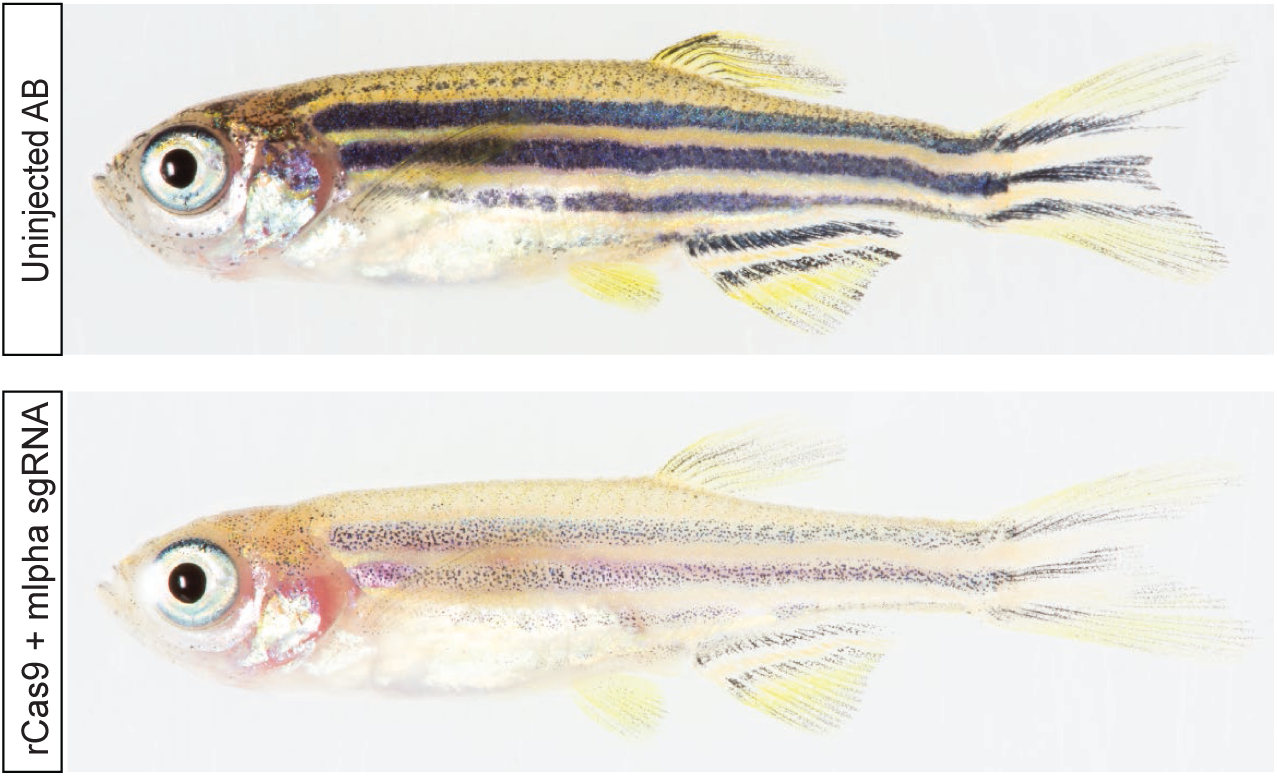
Synthetic sgRNAs produce highly penetrant phenotypes in F_0_ injected animals. Wild-type (AB) embryos were injected with rCas9 and synthetic sgRNA targeting *mlpha*. Uninjected larval animals (top) have normal stripe pattern. Juvenile zebrafish (7 weeks post fertilization) injected with rCas9 and a synthetic sgRNA targeting *mlpha* show reduced pigmentation and phenocopies an established mutant.

**Supplementary Figure 3.**
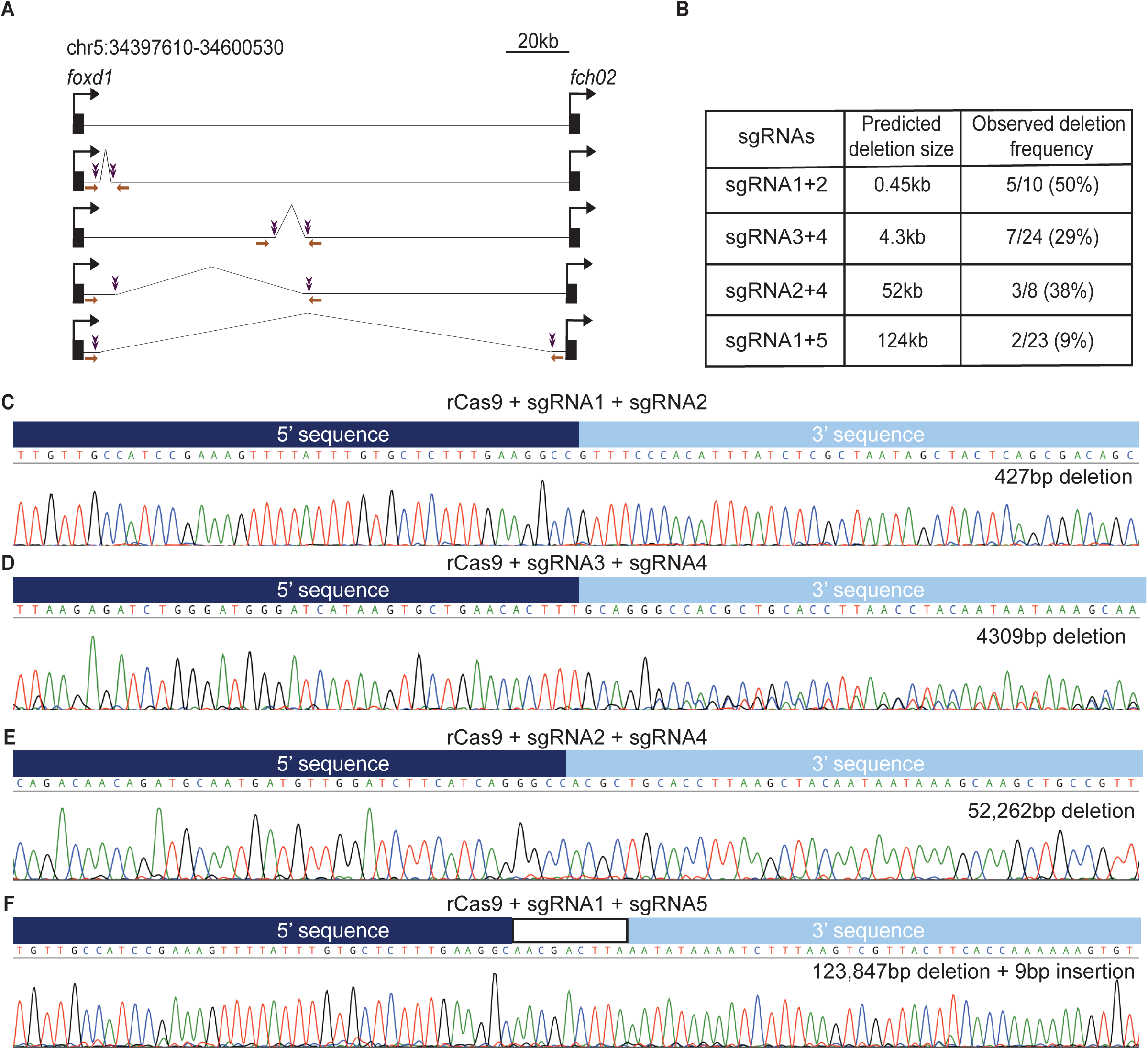
Synthetic sgRNA pairs induce deletions in genomic DNA. (**A**) Schematic of a 126kB genomic region targeted for deletions. Guide locations are marked in purple, and primers used for PCR are displayed as orange arrows. Guide pairs were co-injected into 1-cell stage embryos. (**B**) Table of predicted PCR product size after inducing deletions and observed deletion frequency in individual F_0_ injected embryos. (**C-F**) PCR amplicons from individual embryos were subjected to Sanger sequencing and analysis. The observed deletion size is indicated.

**Supplementary Figure 4.**
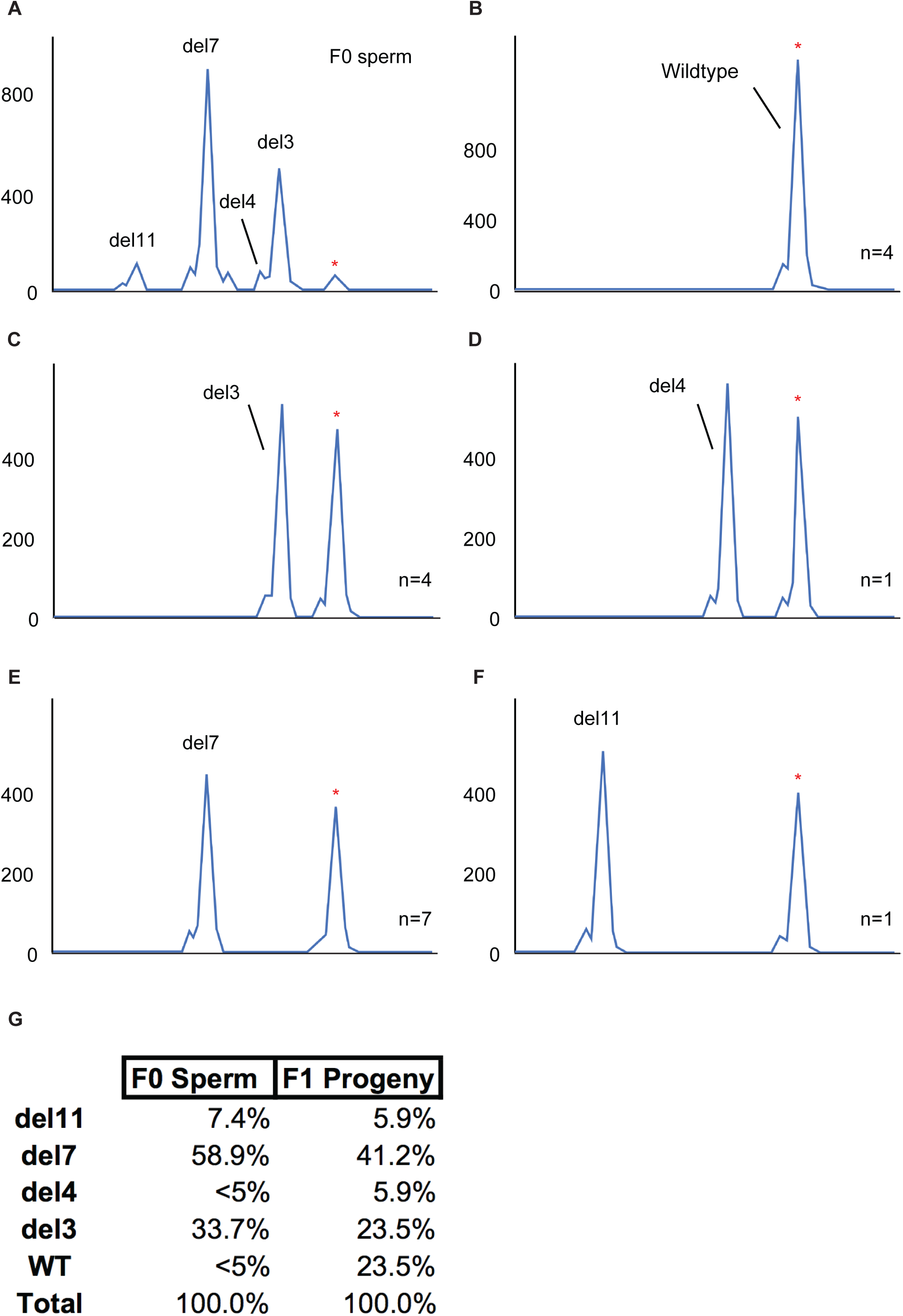
Germline transmission of edited alleles is achieved using synthetic sgRNAs. (**A**) Sperm samples were collected from a CRISPR-injected potential F_0_ founder and assayed using CRISPR-STAT fluorescence-based genotyping. (**B-F**) CRISPR-STAT results for tail clips from representative F_1_ progeny, each carrying one of the distinct alleles with specific insertions and deletions (del3, del4, del7, del11), as well as the wild-type (WT) allele, transmitted from the F_0_ founder. The number of F_1_ offspring identified with each genotype is identified in the bottom right of each panel. Relative fluorescence levels (y-axis) are shown against fragment size in base pairs (x-axis) for each genotyping assay. PCR products consistent with predicted wild type amplicon size are indicated by red asterisks. **(G)** Indels in F_0_ sperm predict alleles transmitted to the germline. Percentages shown are calculated as area under the curve for each peak in the indicated tissue from the F_0_ founder. The frequencies of each mutation in the F_1_ progeny were calculated from a total of 17 siblings from a cross between the F_0_ male parent and a WT female.

**Supplementary Figure 5.**
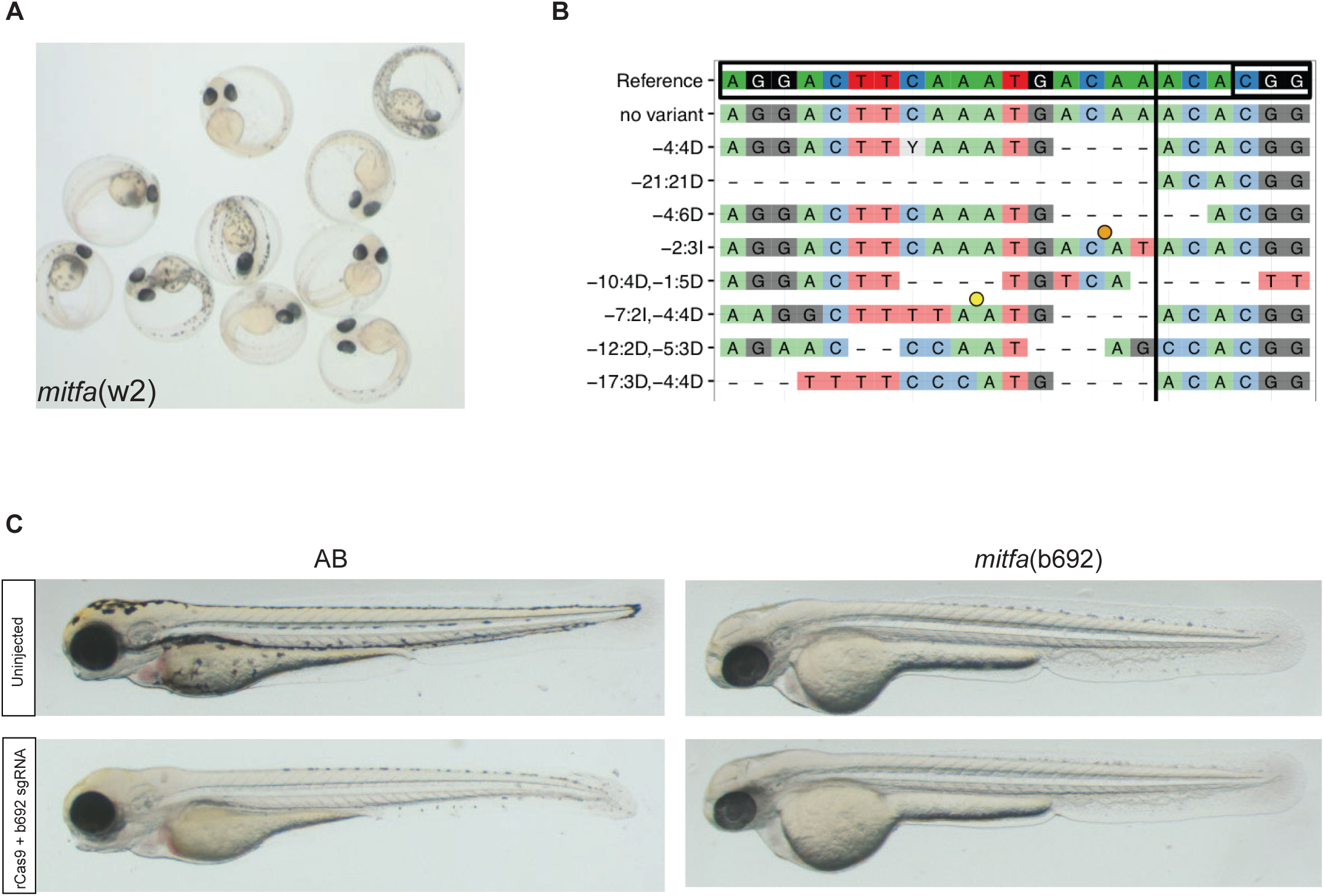
Comparison of melanocyte phenotypes in mitfa mutants after targeting *mitfa* using CRISPR/Cas9. (**A**) *mitfa*(w2) embryos were injected with rCas9 and a specific sgRNA near the mutation site. Ten representative embryos at 24hpf are shown. Melanocyte rescue occurs in a significant fraction of embryos. (**B**) CrispRVariants plot from *mitfa*(w2) embryos injected with rCas9 and nacre gRNA. A majority of embryos harbor indels. (**C**) mitfa(b692) embryos were injected with rCas9 and a synthetic sgRNA near the b692 mutation site (b692 sgRNA). Uninjected wild-type (AB) embryos (top, left) have pigmented melanocytes at 48hpf. Wild-type embryos injected with rCas9 and b692 sgRNA (bottom, left) have a highly penetrant loss of melanocytes. Uninjected *mitfa*(b692) embryos (top, right) lack pigmentation because of the b692 mutation. *mitfa*(b692) embryos injected with rCas9 and the b692 sgRNA (bottom, right) have no evidence of melanocyte rescue.

**Supplementary Table 1.**
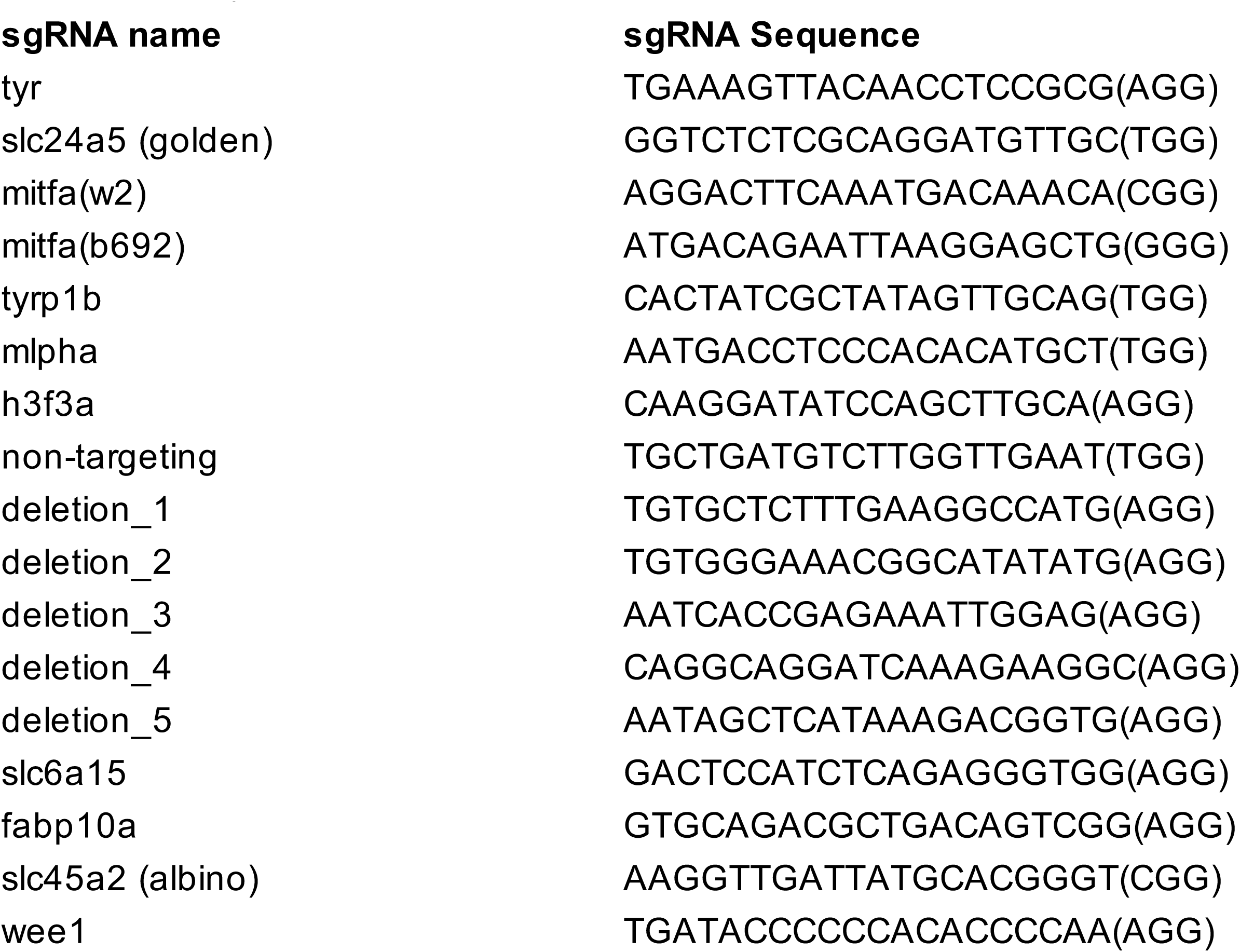
sgRNA sequences. Sequences of sgRNA sequences are noted with the PAM sequence in parentheses.

**Supplementary Table 2.**
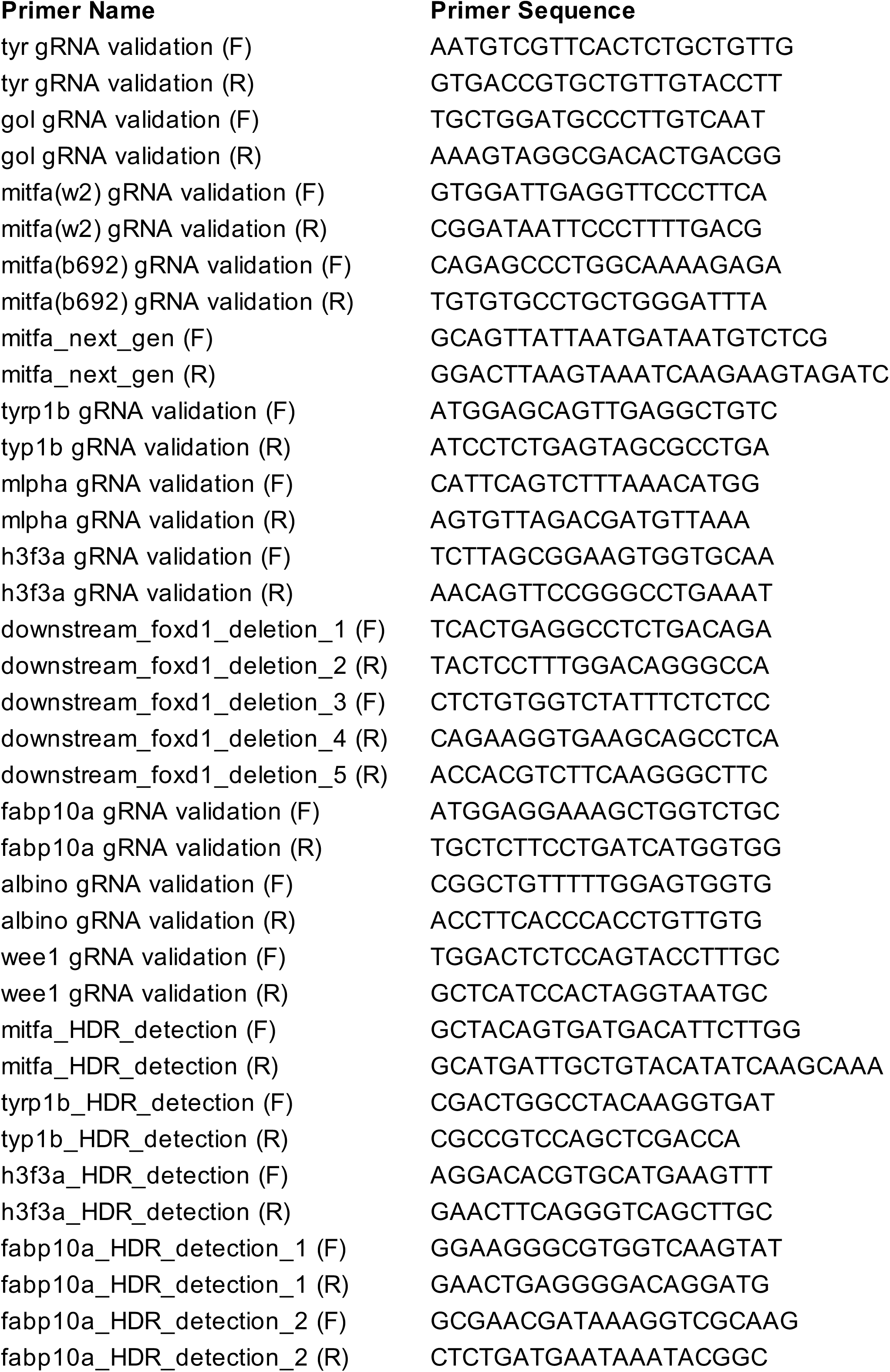
Primer sequences are listed for PCR reactions noted in the text.

